# Testing the gearbox hypothesis for insect flight control

**DOI:** 10.1101/2025.04.29.651276

**Authors:** Abin Ghosh, G. S. Girish Kumar, Sanjay P. Sane

## Abstract

Sustained flight in flies requires wing movements at frequencies ranging between 100–1000 Hz, powered by asynchronous flight muscles whose contraction rate exceeds direct neural stimulation. These muscles attach to the thoracic walls in an indirect flight muscle (IFM) configuration, generating thoracic deformations that drive wing flapping, while smaller steering muscles adjust wing kinematics stroke-by-stroke. The wings articulate with the thorax via the wing hinge, in which the radial stop (RS) and a grooved pleural wing process (PWP) interact to form a mechanical “gearbox” system, proposed to modulate wingbeat amplitude during maneuvering. Supporting this gearbox hypothesis, previous studies observed close RS–PWP interactions during fictive turns in tethered flies. An alternative, though not exclusive, hypothesis suggests that steering muscles govern wingbeat amplitude modulation. However, the gearbox hypothesis had not been rigorously tested in freely flying insects. Here, we first assessed the morphological diversity of the RS–PWP–Pterale C complex across Diptera using scanning electron microscopy and found substantial variation, particularly in PWP morphology. We next tested gearbox function by bilaterally ablating the PWP in freely flying houseflies (*Musca domestica*), which were released into an L-shaped arena requiring 90° yaw turns. Kinematic analyses revealed no significant differences between control and PWP-ablated flies across various wing and body parameters during yaw turns. Both groups could differentially modulate wingbeat amplitude during turns, a key mechanism for yaw torque generation. These findings indicate that the PWP is not essential for wingbeat amplitude modulation, and suggest that steering muscles may primarily control amplitude control during manuevers. Our results challenge the gearbox hypothesis and underscore the flexibility of the dipteran flight control system.

## Introduction

The evolution of flight enabled insects to greatly expand their ecological niches, which led to their spectacular morphological and physiological diversification (Grimaldi & Engel, 2005; Triedel et al, 2024). For instance, insect sizes vary over a range of three orders of magnitude, from a few hundred micrometers in fairy fly wasps (*Kikiki huna*) (Polilov, 2012) to wingspans of 300 mm in the large saturniid moths *Attacus. sp* (Dudley, 2000; Sane, 2016). To generate sufficient lift and remain airborne, smaller insects must compensate for their diminishing wing surface area by increasing their wingbeat frequency. Certain midges (*Forcipomyia sp.*) flap their wings at frequencies exceeding 1000 Hz to remain airborne (Sotavalta, 1953), whereas the common housefly *Musca domestica* flaps its wings at more than 200 Hz. Small insects are thus a good model system to understand how control and coordination are achieved on short timescales, pushing the limits of the ability of the nervous system to acquire and process information and generate rapid motor responses.

Typically, muscles can either contract slowly and generate high force or contract fast and generate low force (Josephson, 1985). To overcome the speed–force trade-off, insects of the orders Diptera, Hymenoptera, and Coleoptera have evolved asynchronous flight muscles that generate high force at high contraction frequencies due to delayed stretch activation and thoracic coupling (Pringle, 1949; Josephson et al., 2000; Deora et al., 2017), enabling high power output for flapping flight. Based on their attachment and orientation within the thorax, the antagonistic flight muscles in the thorax of these insects are classified as the Dorso-Ventral muscles (DVM) which elevate the wings, and Dorsal-Longitudinal muscles (DLM) which depress the wings. Because these muscles attach directly to the thoracic wall, their contractions deform the entire thoracic cavity such that activation of dorso-ventral muscles (DVMs) results in ventral compression and simultaneous longitudinal elongation, which in turn activates the dorso-longitudinal muscles (DLMs) with a delay. Similarly, contraction of DLMs compresses the thorax longitudinally while elongating it dorsoventrally, which activates the DVMs with a delay. This mutual mechanical activation establishes a resonance within the thorax. As a result, these delayed muscle activations can sustain cyclic thorax deformations without the need for neural activation in each cycle (Pringle, 1949; Dickinson and Tu, 1997). These muscles are innervated by motor neurons firing at rates approximately an order of magnitude lower than the wingbeat frequency. The motor input primarily maintains elevated intracellular calcium concentrations, which keeps the muscles in a stretch-activated state and allows for modulation of mechanical power output via changes in spike rate (Gordon and Dickinson, 2006; Lehmann and Bartussek, 2017).

Because the flight muscles of these insects connect to thoracic walls instead of the wing base (Indirect Flight Muscles; IFMs), this arrangement requires a mechanism to transform the deformations of the thoracic box into precisely coordinated upstroke and downstroke movements of the wings. The transmission of thoracic deformations to wing motion is mediated by the wing hinge, a complex assembly of sclerites actuated by muscles. Due to their asynchronous nature, indirect flight muscles (IFMs) cannot provide stroke-to-stroke control. Instead, fine modulation of wing kinematics during maneuvers is achieved by synchronous direct flight muscles (steering muscles), which contract once per neural spike, allowing precise, cycle-specific control. These smaller muscles have well-developed sarcoplasmic reticulum and fast-twitch kinetics (Dickinson et al., 1998; Lehmann and Bartussek, 2017). In *Sarcophaga dux*, 19 such muscles coordinate wing motion, 14 of which insert directly on the sclerites (Deora et al., 2017). While IFMs generate the primary power for wing oscillation, direct muscles refine motion and coordinate left–right wing phasing via a mechanical linkage at the scutellum (Deora et al., 2017). The indirect and direct flight muscles are functionally distinct; whereas indirect muscles generate the primary power required for wing movement, the direct muscles provide fine control for precise adjustments. In addition to power generation and fine control, the coordination of the left-right wing phase is achieved by a mechanical linkage in the thorax called the scutellum (Deora et al., 2015).

To study the control of flight, it is essential to investigate not only the steering muscles but also the mechanics of the sclerites at the wing base, which together form the wing hinge. The wing hinge functions as a system of interconnected levers, enabling forces generated by the muscles to translate into precise wing movements during flight and other behaviours such as courtship (Shirangi et al., 2013). While significant progress has been made in elucidating the combinatorial roles of steering muscles and their effects on wing kinematics (Nachtigall and Wilson, 1967; Bennet-Clark and Ewing, 1968; Wisser and Nachtigall, 1984; Tu and Dickinson, 1994; Trimarchi and Schneiderman, 1995; Fayyazuddin and Dickinson, 1996; Tu and Dickinson, 1996; Heide and Götz, 1996; Dickinson and Tu, 1997; Balint and Dickinson, 2001; Gordon and Dickinson, 2006; Lehmann et al., 2013; Walker et al., 2014; Sadaf et al., 2015; Lehmann and Bartussek, 2017; Lindsay et al., 2017; Melis et al., 2024), the mechanical contribution of the wing hinge itself remains comparatively less understood.

Our understanding of how the wing hinge functions has evolved considerably over the years. Noticing that large flesh fly, *Sarcophaga bullata* under the influence of CCl_4_ anesthesia tended to be in either fully upstroke or fully downstroke positions, Boettiger and Furshpan (1952) proposed a ‘click’ mechanism in Diptera, arguing that the thorax structure was bistable. This meant that the wings adopted one of two stable states at the extreme end of the wing strokes, thus enabling rapid transitions between wing strokes. This view was challenged by Miyan and Ewing (1985), who argued that the ‘click’ was an artefact of the excessive pleuro-sternal muscle tension under CCl₄ exposure based on their observations of wing movements of tethered *Musca domestica, Drosophila melanogaster and Drosophila virilis.* Instead, they proposed that the pleural wing process (PWP) acts as a stopping point and secondary pivot during the downstroke, and its failure to interact with the radial stop (RS) causes abnormal wing movement. Adding to this, Pfau (1987) based on his observation through SEM and observation of the wing hinge of tethered flight *Calliphora erythmcephala* suggested that RS engages differently with the two grooves of PWP, enhancing pronation of the wing and click strength. Ennos (1987) observed RS primarily engaging with the anterior groove of PWP in *Calliphora*, hypothesizing that posterior groove interaction generates asymmetric wingbeat amplitudes for turning. Miyan and Ewing (1988) in tethered flight observations in *Lucilia sericata, Sarcophaga argyrostoma and Glossina morsitans* found instances where RS did not engage PWP, and removing RS led to ipsilateral loss of amplitude control, causing wings to jitter. Wisser (1988) proposed that in *Calliphora erythrocephala* three RS contact points on PWP instead of two, attributing downstroke amplitude variations to these interactions and suggesting PWP is primarily used in extreme maneuvers to avoid structural wear. In addition to the RS-PWP system, traditional descriptions of the wing hinge also include a third component, called the Pterale C (Ritter, 1911), which is thought to function as a mechanosensory structure that provides feedback at wing beat frequency to the wing motor system. Together, the RS-PWP action in conjunction with the function of the Pterale C may be termed as the ‘gearbox’ hypothesis, following the terminology in Nalbach (1989), who conducted extensive kinematic studies on tethered *Calliphora erythrocephala*.

Many aspects of the function of individual gearbox components remain unexplored. For example, scanning electron microscopy (SEM) of the Pterale C reveals a flattened, mushroom-like tip which is covered with very fine bristle-like structures (Miyan and Ewing, 1984). Because the Pterale C is held at a position such that the anterior wing margin collides with it at the end of every downstroke, it is thought to receive stimulation at wingbeat frequency. It has variously been hypothesized to act as a mechanosensor that informs the nervous system about the wing position, or a mechanical damper which ensures smoothness of the wing motion and stroke transitions during flight (Miyan and Ewing, 1984). These hypotheses have also not received experimental confirmation. Miyan and Ewing (1984) recorded sensory activity using extracellular tungsten microelectrodes inserted at the ventral base of the Pterale C, but lacking neural fills from these structures, the site of recording was not clear, nor if Pterale C is indeed a mechanosensory organ. Our own efforts to fill the Pterale C structures in *Sarcophaga spp* were also unsuccessful, especially as there seemed to be no clear nervous connection to the Pterale C (Johri, N., Deora, T, Sane, S.P., Unpublished).

Nalbach (1989) tested the gearbox hypothesis in *Calliphora erythrocephala* using synchronized stroboscopic videography to capture both macroscopic wing motion and microscopic wing hinge mechanics during tethered flight. She identified three distinct downstroke modes based on whether and where the radial stop (RS) contacted the pleural wing process (PWP). These modes correlated with discrete wingbeat amplitudes and were asymmetrically employed during spontaneous turning maneuvers. The study provided experimental evidence that mechanical interactions at the wing hinge could contribute to flight control by modulating stroke amplitude. In studies that examined the role of steering muscles during wing flapping in *Calliphora vicina*, Balint and Dickinson (2001) made similar observations about ‘mode switching’, which they argued was controlled by the muscles of sclerites I and III. In a later study, Deora et al. (2015) observed PWP mode shifts in the flesh fly *Sarcophaga dux* during flight initiation, transitioning from posterior to anterior grooves with increasing amplitude, before RS eventually moved anterior to PWP without interaction (Figure 1A). Walker et al. (2012) showed that in *Eristalis tenax* and *E. pertinax*, the alula—a hinged flap at the wing base—flips between two states during flight, correlating with changes in stroke amplitude, deviation angle, and supination timing. The alula is actuated by the third axillary sclerite and flips at mid-downstroke, matching the timing of RS–PWP engagement. Its flipped state is linked to reduced aerodynamic force and altered body acceleration, suggesting it signals gear shifts. Thus, the alula serves as a visible marker of wing hinge configuration and flight mode changes. Across these studies, PWP emerges as a crucial structure influencing wingbeat amplitude modulation and maneuverability through its interactions with RS. However, all of these studies relied on detailed observations of the wing hinge in tethered flies and its correlation with kinematic parameters to infer the function of the gearbox. Moreover, although these studies were typically carried out on muscomorphic flies, the importance of fine amplitude modulation using gearbox structure extends across all flies, and perhaps also other insects. Hence, a comparative exploration of the gearbox structure is essential in understanding the nuances of the wing hinge and its function in flight. Importantly, it is essential to carry out these studies with freely flying insects, in which multiple strategies of turning may be employed in addition to the gearbox function.

**Figure 1.**
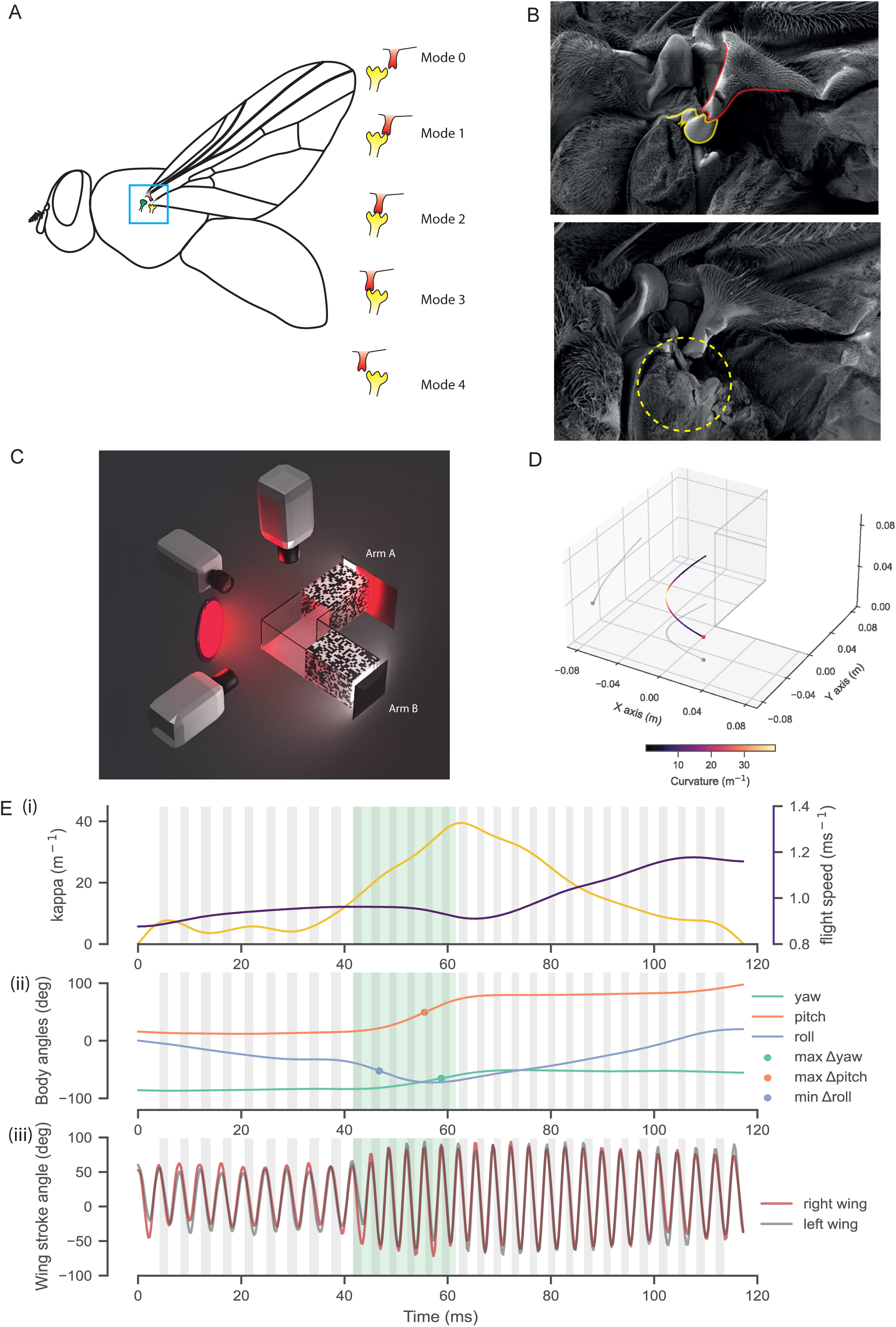
Methodology to investigate the PWP–RS interaction underlying the proposed gear-change mechanism. **A.** Schematic illustrating the possible interaction modes between the radial stop (RS) (red) and the pleural wing process (PWP) (yellow) in *Musca domestica*. In mode 0, the RS is positioned posterior to the PWP without contact. In modes 1–3, the groove on the RS engages sequentially with each of the three peaks of the PWP. In mode 4, the RS moves anterior to the PWP without making contact. **B.** Mechanical ablation of the PWP using microsurgical scissors. (i) The PWP is highlighted in yellow and the RS in red. RS groove is interacting with the middle peak of the PWP. (ii) The dotted yellow circle indicates the site of ablation following removal of the PWP. **C.** L-shaped flight arena used to induce left turns in Musca domestica. Flies initiated flight from arm A toward arm B in an L-box covered with a high-contrast checkerboard pattern for visual guidance. Three high-speed cameras were placed approximately orthogonal to each other to capture top, front, and side views at the junction for 3D reconstruction of flight trajectories. The arena was illuminated with high-intensity tungsten lighting filtered through a deep red filter to provide sufficient illumination for high-speed recording while minimizing behavioral interference. White LED panels positioned at the ends of arms A and B acted as visual attractants to encourage movement between arms. Once flies settled in arm A, a red filter was slid across the backlight, inducing a startle response through a sudden perceived decrease in light intensity and causing the flies to fly toward the junction and turn left into arm B. Two additional white LED panels (not shown) were positioned above the checkered walls to increase ambient light levels and further support sustained flight. **D.** Center of mass trajectory of a flying Musca domestica. The trajectory is color-coded by local curvature (κ), computed using the Serret–Frenet framework, where κ quantifies the instantaneous turning radius along the flight path. **E.** Overview of analyzed flight data. The green shaded region highlights the segment selected for further analysis based on centripetal force (κv²) maxima. The grey shaded regions indicate periods of wing downstroke. (i) Curvature (κ) (yellow) and speed (violet) of Center of mass. (ii) Body angles (yaw (green), pitch (orange), roll (blue) ), with dots marking the points of maximum rate of change for each rotational axis. (iii) Wing stroke angles for the left (dark grey) and right wings(red).

The gearbox hypothesis is predicated on the assumption that the RS-PWP interaction is crucial for amplitude modulation. An alternative hypothesis is that wing amplitude is primarily modulated by the steering muscle control of the sclerites. These two hypotheses are not mutually exclusive, however, as both the wing hinge and the steering muscles are essential components of the wing transmission system, that converts thoracic vibrations to wing movement. Testing these hypotheses requires us to address several questions. First, how variable is the gearbox morphology across Diptera? Second, how do the relative configurations of RS and PWP influence the wing and body movements during turns? Third, how does selective modification of the wing hinge structure compromise the flight of insects? To address these questions, we investigated the structural and functional role of the wing hinge in flight control, focusing on the RS-PWP interaction. We conducted an electron microscopic examination of the wing hinge, focusing on PWP, RS and Pterale C morphology across several members of the Diptera to determine the variability in their structures. We next conducted a focused investigation of the functional role of the RS-PWP system in modulating wingbeat amplitude in freely flying houseflies *Musca domestica*.

## Methods

### Scanning Electron Microscopy (SEM) of the wing hinge

Wild dipteran insects were collected opportunistically from the campus of NCBS, Bangalore, using soft insect nets. After collection, they were fixed in 4% paraformaldehyde (PFA) to preserve the structural integrity of their thorax and associated structures. Wings were positioned in the fully elevated position during fixation to facilitate imaging of the wing hinge. Fixed specimens underwent serial dehydration in ethanol grades of 20%, 30%, 50%, 60%, 75%, 90%, and twice in 100%, for 10 minutes each. To prevent morphological distortion during drying, the ethanol-dehydrated samples were subjected to critical point drying using a Leica EM CPD 300 Critical Point Dryer (Leica Microsystems, Wetzlar, Germany) for 90 minutes. Dried samples were mounted on aluminium stubs using conductive carbon adhesive tape. A gold coating (∼10 nm) was applied using an Emitech K550X sputter coater (Quorum Technologies Ltd., West Sussex, UK) for 5 minutes to enhance conductivity. Imaging was performed using a Zeiss Merlin Compact VP SEM (Zeiss, Germany) at an accelerating voltage of 2–5 kV and a working distance of 10–15 mm using the SE2 detector.

### Animal Maintenance for free flight experiments

Adult *Musca domestica* specimens were collected from wild populations and housed in ventilated containers with *ad libitum* food and water using cotton pads soaked in a 10% sugar solution and freshwater, at 25 ^O^C. Specimens were used within two days of collection to ensure physiological consistency.

### Pleural Wing Process Ablation

Male *M. domestica* specimens with no wing damage, mites, or other visible deformities were selected from the housing container for PWP ablation. Flies were anesthetized in a 50 mL centrifuge tube and kept on ice for 1 minute. To maintain anaesthesia during the ablation procedure, each specimen was positioned on a brass block maintained on ice. To expose the PWP, the wings were held in the upstroke position and the PWP was bilaterally ablated using fine microscissors (15000-08, Fine Science Tools, California, USA) (Figure 1B). The ablation procedure, including anesthetization, was completed within five minutes to prevent any adverse effects of prolonged exposure to ice. Flies were allowed to recover in a 50 mL centrifuge tube overnight with *ad libitum* water but no sugar. Any specimens exhibiting unusual wing resting positions after overnight recovery were excluded from further experimentation to ensure consistency in the data. Control specimens underwent the same anesthetization protocol and time but did not undergo PWP ablation.

### Free Flight Experimental Setup to Induce Left Turns in Houseflies

Free flight experiments were conducted in an L-shaped flight box designed to induce a left turn during flight, starting from arm A towards arm B (Figure 1C). The arms of the box, except at the junction, were covered with a high-contrast black-and-white random checkerboard pattern to provide visual cues for navigation. Three high-speed cameras v1212, VEO 640L, and v611 (Phantom, Vision Research Inc.), were set up roughly orthogonal to each other at the junction to capture top, front, and side views, enabling 3D reconstruction of flight trajectories. The cameras operated at 4000 frames per second (FPS) with an exposure time of 70 microseconds to obtain a crisp image of their wing outlines. The flight box was illuminated from the top using a white LED panel to provide adequate lighting for flight (∼ 3500 lux). A bright tungsten lamp covered with a red filter provided additional illumination optimized for the cameras without interfering with the flies’ responses to visual stimuli. At the back end of arm A, a white LED panel was used to trigger a startle response in the flies. When a red filter was suddenly slid in front of this light, the flies exhibited a reflexive startle response and flew toward the junction of the L-box. Upon reaching the junction, the flies turned left and proceeded toward the white light at the end of arm B (Figure 1C). The pleural wing process ablation was verified after the experiment using a dissection microscope to ensure the integrity of the ablation procedure.

### Calibration

The flight box was securely mounted on an opto-mechanical breadboard equipped with multiple guide bolts to maintain a consistent position relative to the cameras throughout the experiments. Calibration of the three camera views (top, front, and side) was performed based on easyWand (Theriaul et al. 2014). A thin capillary of 90 mm, suspended by a thread, was moved around the junction of the L-box in various orientations and positions. The digitized points from the end of the capillary are used to estimate 11 parameters of the Direct Linear Transformation (DLT) matrix, which maps the 3-D projection of the real-world point onto the three 2D camera views. The 3D coordinate system was defined with the lower inner corner of the L-box at the junction as the origin. The arm A of the box was aligned with the y-axis, the arm B was aligned with the x-axis, and the height of the box was aligned with the z-axis. The DLT coefficients are updated to add the rotational and translational components to meet the above criterion. This coordinate system provides a consistent frame of reference for analyzing flight kinematics across trials.

### Digitization of the Video and Analysis

To quantify the flight kinematics, six key anatomical points - the head, left-wing base, left-wing tip, right-wing base, right-wing tip, and abdomen tip - were manually annotated for each camera view using DLTdv8 (Hedrick, 2008). The 3D coordinates of these points were calculated using the DLT calibration coefficients derived from the wand calibration method.

For body kinematic analysis, the 3D coordinates were low-pass filtered using a Butterworth filter with a cut-off frequency of 100 Hz to remove high-frequency oscillations from wing oscillations close to 200 Hz while preserving body motion characteristics. For wing kinematics analysis, the coordinate points were low-pass filtered using a Butterworth filter with a cut-off frequency of 1000 Hz to remove high-frequency noise.

### Flight Speed and Curvature of Flight Trajectory

The center of mass (COM) of the fly was estimated as the midpoint between the left and right-wing bases. The velocity vector (v) was derived as the first temporal derivative of COM position. To calculate the curvature **κ** of the fly’s COM trajectory in 3D, we used the Frenet– Serret frame. The Frenet–Serret frame describes the local geometry of a curve in space using three mutually orthogonal unit vectors: the tangent T (direction of motion), the normal N (direction of turning), and the binormal B (orthogonal to both).

The rate of change of the tangent vector T with respect to arc length s is directed along the normal vector N and scaled by the curvature κ:

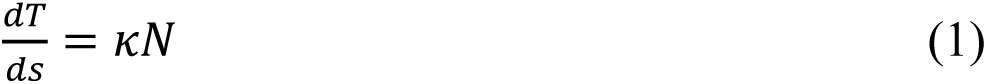

We used a custom MATLAB function, which computed the Frenet-Serret frame of a 3D curve. Curvature was computed from the three-dimensional center of mass trajectory recorded as time series data in x, y, and z coordinates. The first step involved estimating the velocity vector at each time point by numerically differentiating the position data using finite differences. The speed, or arc length differential, was then calculated as the magnitude of the velocity vector. This speed was used to normalize the velocity and obtain the unit tangent vector, which describes the direction of motion along the trajectory. Next, the rate of change of the tangent vector with respect to time was computed, and this derivative was rescaled using the arc length to obtain the change of tangent vector with respect to distance travelled along the curve. Curvature was finally calculated as the magnitude of this arc-length-based derivative of the tangent vector (Figure 1D).

### Calculation of Dynamic Body Angles

Dynamic changes in yaw, pitch, and roll were computed iteratively for each frame based on the head and wing base vectors. Initial values of yaw, pitch, and roll with respect to the global coordinate system were determined using the head and wing base vectors from the first frame. Yaw and pitch were obtained by converting the head vector into spherical coordinates, while roll was calculated from the transformed base vector.

For each subsequent frame, the local coordinate system was established using the following basis vectors: the z-axis as the unit vector of the cross-product of the head and base vectors, the x-axis as the normalized head vector, and the y-axis as the cross-product of the z-axis and x-axis. The transformed base vector was then rotated to align with the yaw and pitch adjustments, and roll was computed from its spherical coordinates. The head and base vectors of the current frame were transformed into the defined local coordinate system. Changes in yaw and pitch were then calculated by converting the transformed head vector into spherical coordinates. The cumulative sums of *Δ* yaw, *Δ* pitch, and *Δ* roll were added to their respective initial values to compute the overall yaw, pitch, and roll for each frame (Figure 1E).

### Calculation of wing parameters

To calculate the wing angles based on the stroke plane, we used a custom MATLAB function that processes the 3D spatial coordinates of the wing base and tips across time. The method calculated the vectors representing each wing and rotated these vectors to align with the body-centric yaw, pitch, and roll. The wing angles were then decomposed into azimuth (stroke angle), stroke deviation, and wing length in spherical coordinates. The stroke plane angles were computed by determining the orientation of the plane formed by the wing vectors and the body plane, accounting for variations in wingbeats. These angles were further refined by aligning the wing motion with the averaged stroke plane vector for an accurate representation of wing kinematics. This approach enabled precise quantification of wing motion dynamics relative to the stroke plane while the insect is freely flying (Figure 1E).

## Results

### The grooved gearbox morphology is not conserved across Dipterans

The gearbox hypothesis is predicated on the assumption that the RS-PWP interaction is crucial for amplitude modulation, and hence must be conserved across Diptera. We compared the RS, PWP and Pterale-C structures in the wing hinge in ten members sampled across the Dipteran phylogenetic tree. These included *Sarcophaga dux*, *Calliphora spp.*, *Musca domestica*, *Drosphila melanogaster, Tabanus spp., Hermetia illucens, Clogmia spp., Pselliophora spp., Allobaccha spp.,* Dolichopodidae (*genus indetermined)*. The Radial Stop (RS) was identified as the protrusion from the radial vein of the wing at the base and the Pleural Wing Process (PWP) as the protrusion near the base of the wing arising from the pleural wall. The Pterale C structure, when present, is also clearly identifiable as the third component of the wing hinge, arising from the thoracic wall anterior to PWP. In the Dipterans sampled here, Pterale C was absent in three fly species, including Psychodomorpha (drain flies), Tabanomorpha (horseflies), and Empidoidea (long-legged flies), which are not monophyletic. This suggests that Pterale C may be a conserved feature of all Diptera, but secondarily lost in these three members.

RS and PWP were present and identifiable across all ten specimens. However, their morphology is quite variable, in a manner that does not seem phylogenetically related. For example, in the groups sampled here, the structure of the RS is ungrooved in the Dipteran groups Tipulomorpha, Psychodomorpha, Stratiomyomorpha, and Platypezoidae. In Empidoidea and Ephydroidea, the PWP contains a single groove. In Tabanomorpha and Oestroidae, the PWP is multi-grooved (Figure 2 A-J).

**Figure 2.**
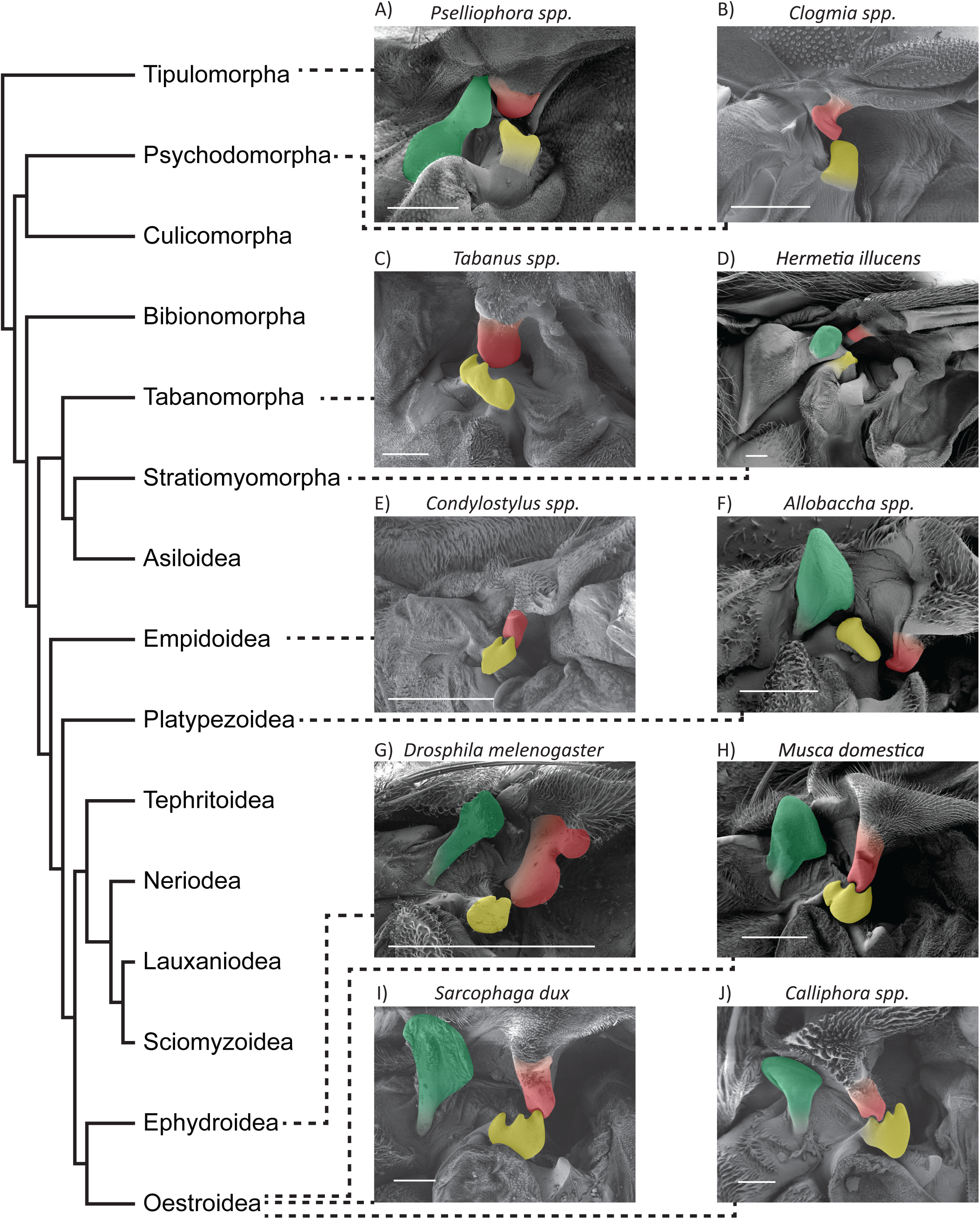
Morphological diversity of the pleural wing process (PWP) and radial stop (RS) across Diptera. Scanning electron micrographs of the wing bases of representative dipteran species, sampled across the dipteran phylogeny. The Pterale C is false-colored in green, the pleural wing process (PWP) in yellow, and the radial stop (RS) in red. Species were selected to capture morphological variation across major dipteran lineages. The phylogenetic relationships among the sampled species are shown alongside (phylogenetic tree reproduced from Wiegmann et al., 2011).

These findings demonstrate that the two-grooved morphology of the PWP is not conserved across Diptera, indicating significant structural divergence. Although detailed and standardized flight performance measurements in these groups are not available, cursorily, there appears to be no correlation between flight ability and wing hinge structure in any of these flies. For example, horseflies (Tabanomorpha) are known to fly at high speeds and long distances (Wilkerson and Butler, 1984) but lack the Pterale C. Similarly, hoverflies (Platypezoidea) are extremely maneuverable, which requires exquisite modulation of wing amplitude (Walker et at, 2010), but their PWP is ungrooved. In the case of Tabanomorpha and Oestroidea, the RS interdigitates with PWP grooves, which strongly hints that their shapes have evolved to interact and minimize slippage during such interactions. Together, these data suggest that the gearbox morphology is highly variable and its structure does not clearly correlate with flight ability, as would be predicted by the gearbox hypothesis.

### The flies performed the left turn through a banked turn

How important is the PWP for generating free-flight turns? The gearbox hypothesis posits that the PWP is an essential component in wingbeat amplitude modulation, without which aerial yaw turns are not possible. Hence, we next analysed free-flight kinematics of houseflies as they performed yaw turns in an L-shaped flight chamber. The yaw maneuver requires the flies to bank and consists of three distinct phases. First, a roll phase in which a leftward roll orients their dorsal side toward the inner side of the turn. Second, a pitch phase, in which they pitch up to complete the turn. Third, a counter-roll phase, in which they return to a dorsal-side-up orientation after completing the turn (Movie S1). Of the 20 flies, 18 consistently exhibited these distinct phases. The two exceptions had already oriented with their dorsal side toward the inner turn before entering the calibrated volume, thus the first roll phase was not observed. These results demonstrate that flies employ a structured kinematic strategy when performing directional turns. We next compared these kinematic patterns in 20 houseflies, including 10 control individuals and 10 flies with bilaterally-ablated PWP.

### Whole body kinematic parameters of control and PWP-ablated flies are comparable

We first analysed the flight speed and curvature of the flight trajectories to compare the flight performance between the control and PWP-ablated groups. The group median of median speed during the maneuver for the control group was 0.99 ms^-1^, not significantly different from the group median of median speed of the PWP-ablated group 0.97 ms^-1^ (Figure 3B; n=10, Mann–Whitney U rank test, p=0.850). Similarly, the median value of the group median of maximum speed achieved by the PWP-ablated flies 1.05 ms^-1^ was comparable to the group median of maximum speed of the control group 0.97 ms^-1^ (Figure 3A; n=10, Mann–Whitney U rank test, p=0.734).

**Figure 3.**
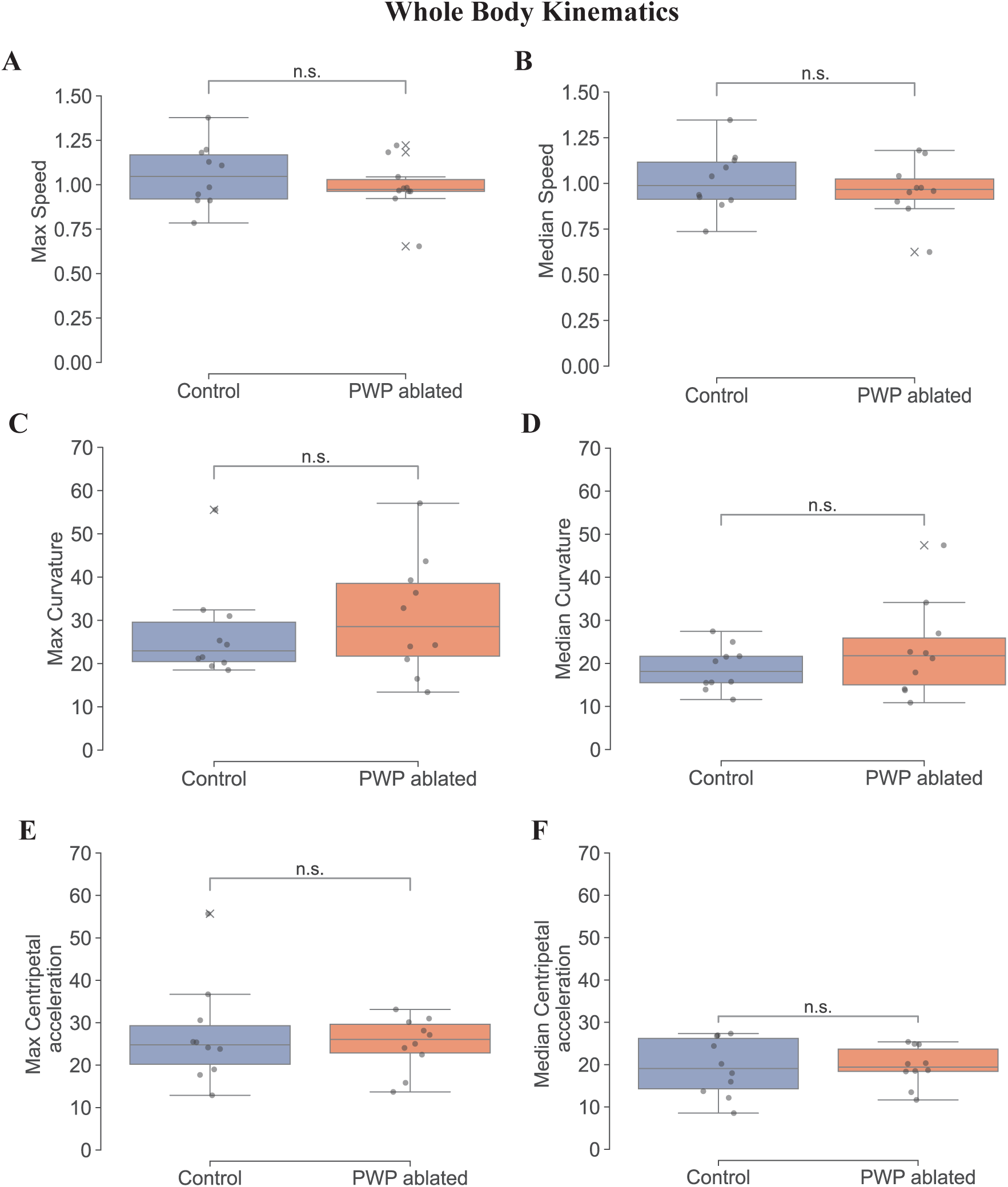
Whole body kinematic parameters of control and PWP-ablated flies. **A**. Maximum flight speed during the maneuver was not significantly different between control and PWP-ablated groups (Mann–Whitney U test, p=0.734; n=10 per group). **B.** Median flight speed during the maneuver was also not significantly different between groups (p=0.850). **C.** Maximum curvature (κ) showed no significant difference between control and PWP-ablated groups (p=0.521). **D.** Median curvature (κ) was similarly not significantly different between groups (p=0.521). **E.** Maximum centripetal acceleration (κv²) did not differ significantly between the groups (p=0.970). **F.** Median centripetal acceleration (κv²) was also statistically indistinguishable between control and PWP-ablated flies (p=0.970).

The median value of the group median of median curvature observed in the control flies was 18.12 m^-1^, which again did not differ significantly from that in PWP-ablated flies 21.79 m^-1^ (Figure 3D; n=10, Mann–Whitney U rank test, p=0.521). The group median of maximum curvature achieved by the PWP-ablated group 22.94 m^-1^ was also statistically indistinguishable from the of the control flies 28.56 m^-1^ (Figure 3C; n=10, Mann–Whitney U rank test, p=0.521).

We next analysed centripetal acceleration (κv^2^), which was also not significantly different between the two groups. The median of control group’s median centripetal acceleration was 19.08 m s^-2^, comparable to the PWP-ablated group 19.42 m s^-2^ (Figure 3F; n=10, Mann– Whitney U rank test, p=0.970). Likewise, the group median of the maximum centripetal acceleration achieved by the PWP-ablated flies 24.78 m s^-2^ was not significantly different from that of the control group 26.08 m s^-2^ (Figure 3E; n=10, Mann–Whitney U rank test, p=0.970).

These results indicate that PWP ablation does not significantly alter flight performance as measured by speed, curvature, or centripetal acceleration.

### Analysis of roll, pitch and yaw body angles during turning maneuvers shows no significant difference

To assess differences in flight dynamics between control and PWP-ablated flies, we calculated the rates of change of the body angles—pitch, roll, and yaw. These analyses again revealed no significant difference between the two groups for any of the parameters.

The rate of change of yaw for the control group median of the median value of 1458.32°s^-1^, which was not significantly different from the median value of 1513.99°s^-1^ observed in the PWP-ablated group (Figure 4D; n=10, Mann–Whitney U rank test, p=0.910). Similarly, the group median of the maximum yaw rate in the control group 2273.04°s^-1^ was comparable to the PWP-ablated group 2004.62°s^-1^ (Figure 4C; n=10, Mann–Whitney U rank test, p=0.345).

**Figure 4.**
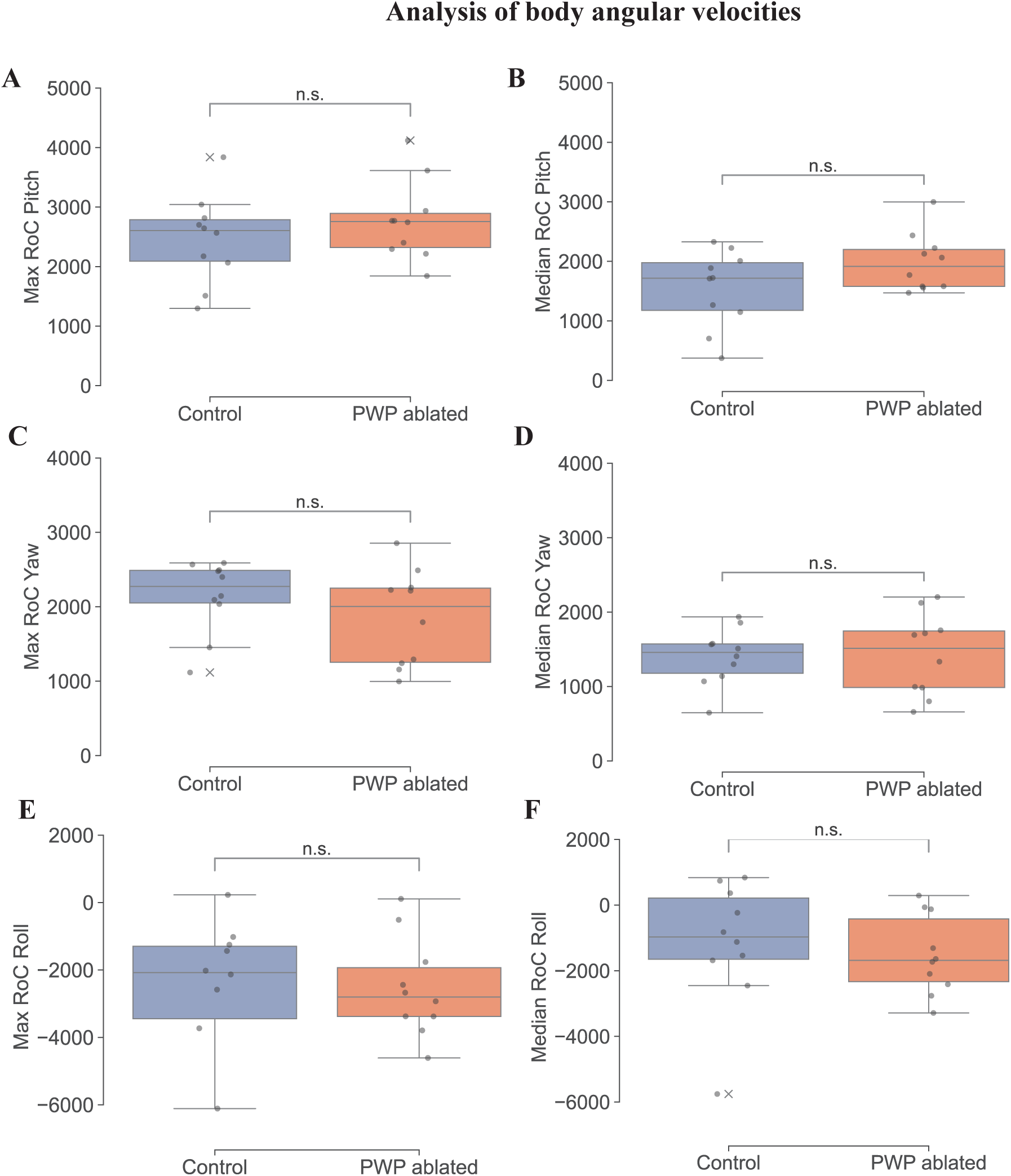
Analysis of body angular velocities (yaw, pitch, and roll) during turning maneuvers in control and PWP-ablated flies. **A.** Maximum pitch rate showed no significant difference between control and PWP-ablated groups (Mann–Whitney U test, p=0.385; n=10 per group). **B.** Median pitch rate was also not significantly different between groups (p=0.241). **C.** Maximum yaw rate was statistically comparable between control and PWP-ablated groups (p=0.345). **D.** Median yaw rate showed no significant difference between groups (p=0.910). **E.** Maximum roll rate along the negative axis did not differ significantly between groups (p=0.678). **F.** Median roll rate along the negative axis was also not significantly different between control and PWP-ablated flies (p=0.307).

For pitch, the median rate of change observed in the control group was 1716.53°s^-1^, and this was statistically similar to the median pitch rate in the PWP-ablated group 1916.95°s^-1^ (Figure 4B; n=10, Mann–Whitney U rank test, p=0.241). The median value of the maximum pitch rate achieved by the PWP-ablated flies 2605.82°s^-1^ was also not significantly different from that of the control group 2757.33°s^-1^ (Figure 4A; n=10, Mann–Whitney U rank test, p=0.385).

Analysis of roll revealed that the median roll rate for the control flies -972.25°s^-1^ was indistinguishable from the corresponding value in the PWP-ablated flies -1684.43°s^-1^ (Figure 4F; n=10, Mann–Whitney U rank test, p=0.307). Additionally, the median of the maximum roll rate in the negative axis for the control group -2076.96°s^-1^ did not significantly differ from that of the PWP-ablated group -2801.06°s^-1^ (Figure 4E; n=10, Mann–Whitney U rank test, p=0.678).

Thus, PWP ablation did not affect the rate of change of pitch, roll, or yaw during flight, suggesting that body angle dynamics are robust to the loss of the PWP.

### Flies with PWP ablated can modulate wing beat amplitude

The group median of the median wingbeat frequencies during the turn in the control group was 267 Hz, which was not significantly different from the group median value observed in the PWP-ablated group, 276 Hz (n=10, Mann–Whitney U rank test, p=0.413; Figure 5B).

**Figure 5.**
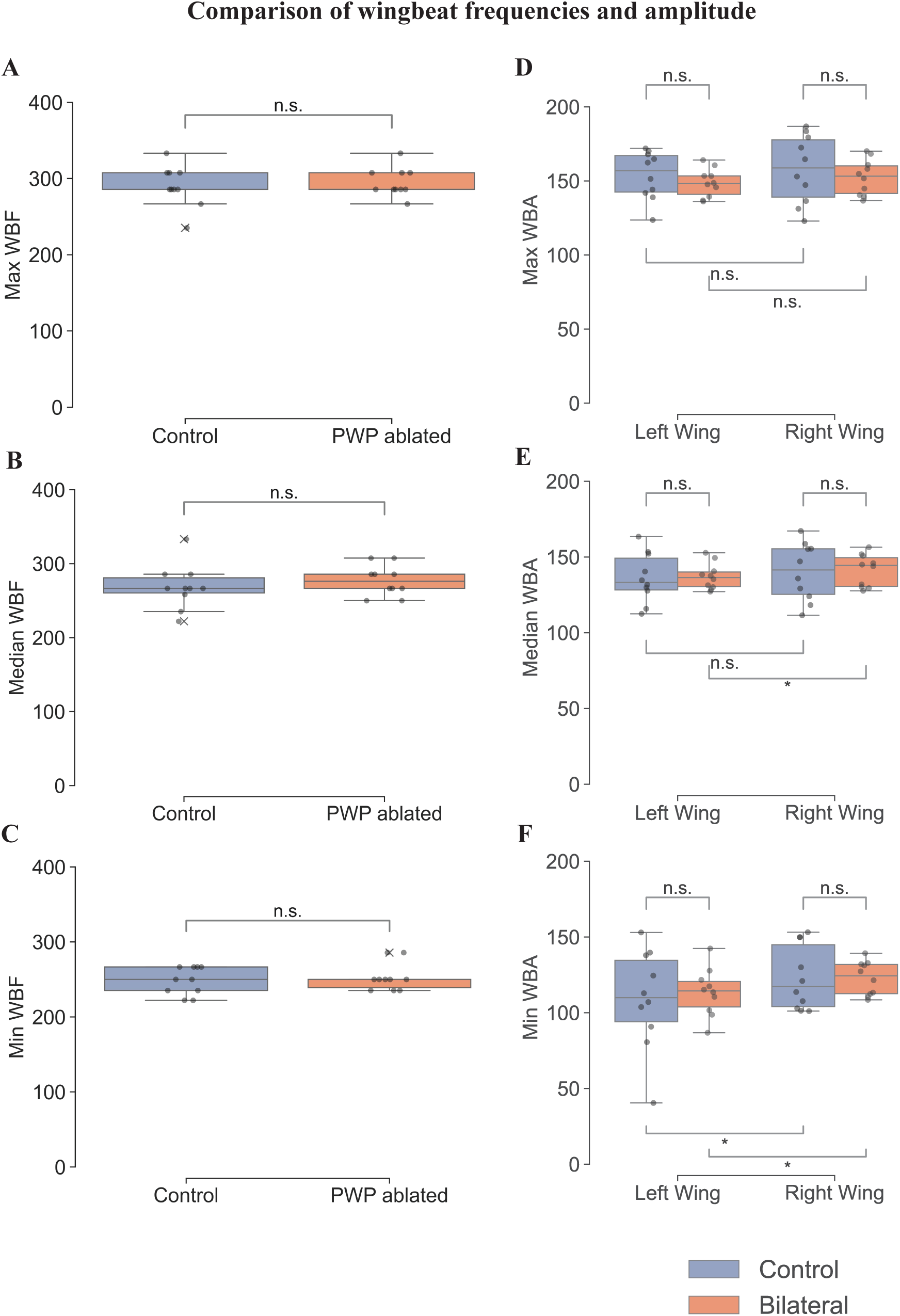
Comparison of wingbeat frequency and wingbeat amplitude between control and PWP-ablated flies during leftward turning maneuvers. **A.** Maximum wingbeat frequency was not significantly different between control and PWP-ablated groups (Mann–Whitney U test, p=0.809; n=10 per group). **B.** Median wingbeat frequency showed no significant difference between groups (p=0.413). **C.** Minimum wingbeat frequency was also statistically similar between groups (p=0.809). **D.** Maximum wingbeat amplitude comparisons showed no significant differences between left and right wings within control or PWP-ablated groups, and no significant differences between groups for either wing. **E.** Median wingbeat amplitude was not significantly different between left and right wings in control flies (p=1.000), but showed a significant asymmetry in PWP-ablated flies (p=0.014); however, no differences were found between groups when comparing left or right wings separately. **F.** Minimum wingbeat amplitude was significantly lower in the left wing compared to the right wing in both control (p=0.020) and PWP-ablated flies (p=0.027); no significant differences were found between groups for either wing.

Similarly, the group median of the maximum wingbeat frequencies was 285 Hz for both the control and PWP-ablated flies, with no significant difference between the groups (n=10, Mann–Whitney U rank test, p=0.809; Figure 5A). In addition, the group median of the minimum wingbeat frequencies was 250 Hz in both the control and PWP-ablated groups, with no significant difference detected (n=10, Mann–Whitney U rank test, p=0.809; Figure 5C).

When comparing the group medians of the minimum wing beat amplitudes during a left turn, the left wing showed significantly lower amplitudes than the right wing in both control flies (n=10, Mann–Whitney U rank test, p=0.020; Figure 5F) and PWP-ablated flies (n=10, Mann– Whitney U rank test, p=0.027; Figure 5F), suggesting that flies reduce the wingbeat amplitude of the left wing to execute a leftward turn. However, when comparing the group medians of the minimum wing beat amplitude of the left wing between control and PWP-ablated groups, no significant difference was observed (n=10, Mann–Whitney U rank test, p=0.850; Figure 5F), and similarly, no significant difference was found for the right wing between the two groups (n=10, Mann–Whitney U rank test, p=0.678; Figure 5F).

In contrast, when comparing the group medians of the maximum wingbeat amplitudes, no significant differences were detected between the left and right wings within the control group (n=10, Mann–Whitney U rank test, p=0.557; Figure 5D) or within the PWP-ablated group (n=10, Mann–Whitney U rank test, p=0.084; Figure 5D). Moreover, comparison of the group medians of the maximum amplitude of the left wing between control and PWP-ablated flies revealed no significant difference (n=10, Mann–Whitney U rank test, p=0.345; Figure 5D), and the right wings also did not differ significantly between groups (n=10, Mann– Whitney U rank test, p=0.623; Figure 5D).

For the group medians of the median wingbeat amplitudes, no significant difference between the left and right wings was found in control flies (n=10, Mann–Whitney U rank test, p=1.000; Figure 5E). However, in PWP-ablated flies, the group median amplitude of the left wing was significantly lower than that of the right wing (n=10, Mann–Whitney U rank test, p=0.014; Figure 5E), indicating an asymmetry introduced after ablation. Despite this within-group difference, the group median amplitude of the left wing did not differ significantly between control and PWP-ablated groups (n=10, Mann–Whitney U rank test, p=0.850; Figure 5E), nor was there a significant difference in the right wing group medians between the groups (n=10, Mann–Whitney U rank test, p=1.000; Figure 5E).

Together, these results show that flies turn by selectively reducing the minimum wing beat amplitude of the inner (left) wing, a mechanism that remains functional after PWP ablation. Maximum wingbeat amplitude was not different between the left and right wings or between the groups, indicating it is not modulated during a turn. A difference in median wingbeat amplitude emerged only after ablation, suggesting minor shifts in baseline wing kinematics. Overall, the ability to asymmetrically adjust minimum wingbeat amplitude during turning is preserved despite PWP removal.

## Discussion

### How does a fly generate a yaw turn?

To initiate a yaw turn, flies typically create asymmetric stroke amplitudes between the left and right wings, generating net torque around the vertical axis (Fry et al., 2003). Beyond amplitude differences, precise control of yaw is further refined through modulation of wing rotation during stroke reversals, which alters the relative aerodynamic angle of attack (Muijres et al., 2014). Additionally, yaw turns are often accompanied by a banked maneuver, in which flies produce coordinated pitch and roll torques via fine-tuned adjustments to multiple wing kinematic parameters. These banked evasive maneuvers enable much faster reorientation, achieving angular velocities up to 5300° s⁻¹, compared to approximately 1000° s⁻¹ during voluntary yaw turns (Muijres et al., 2014). During such maneuvers, the body rolls in the direction of the turn, redirecting aerodynamic forces laterally to facilitate smooth changes in heading (Dickinson & Muijres, 2016).

In the experiments described here, flies were required to perform yaw turns to change their flight direction (Fig 1 A-E). Because pitch and roll are inherently coupled during banked yaw maneuvers, this behavior offered an ideal context in which to examine the subtle impairments that may result from PWP ablation. Our findings indicate that both control and PWP-ablated flies executed turns using a banked maneuver. Thus, their ability to coordinate pitch, roll, and yaw components during flight was unimpaired by PWP ablation, which in turn suggests that the interaction between RS and PWP may not be critical in generating complex maneuvers.

Alternatively, in absence of RS-PWP interactions, flies may rely on activity from steering muscles to generate the necessary changes in kinematics to generate complex manuevers.

### Role of the gearbox in Dipteran flight

The gearbox mechanism has also been identified in other insect orders, where it exhibits considerable variation despite the ability of these insects to smoothly modulate their wing amplitude. In flies, the interaction between the radial stop (RS) and the multi-grooved pleural wing process (PWP) is believed to facilitate unilateral amplitude modulation of each wing. As observed in *Sarcophaga dux* and other flies (Deora et al., 2015), the RS transiently contacts one of the PWP grooves during the downstroke, serving as a stop that constrains wingbeat amplitude to one of five modes. Among these, mode 0 corresponds to the disengaged state, where the wing remains decoupled from thoracic vibrations. In contrast, modes 1–3 involve engagement of the wing clutch, each associated with a specific wingbeat amplitude (Nalbach, 1989; Deora et al, 2015). In modes 1, 2 and 3, the RS contacts the peaks 1, 2 and 3 of the PWP, respectively, whereas in mode 4, the RS moves anteriorly without making contact. Consistent with this observation, Nalbach’s (1989) tethered flight observations described three modes based on amplitude distributions.

The gear change mechanism, as described above, requires precise actuation within a 10 µm range at wingbeat frequencies exceeding 100 Hz. Achieving such fine control would in turn require rapid sensory feedback about the wing hinge configuration, but it is not clear what would be the source of this feedback. Furthermore, as shown here, the number of PWP grooves varies substantially among fly species with flight frequencies above 100 Hz, yet all maintain continuous amplitude modulation. Further, our SEM analyses across Oestroidea species (*Calliphora spp*., *Sarcophaga dux*, and *Musca domestica*) revealed that the RS itself possesses a distinct groove and interlocks precisely with the peak of the PWP. Note that Mode 1,2 and 3 are separately captured in Fig 2I, H, and J respectively, *albeit* in different Oestroidea species.

The experiments described here explicitly test the role of the gearbox in wing amplitude modulation during aerial maneuvers. Insects adjust wingbeat amplitude to generate asymmetrical forces for steering or to counteract deviations from stable flight caused by external disturbances. If the pleural wing process (PWP) plays a critical role in wingbeat amplitude modulation, its bilateral ablation should significantly impact flight performance in the housefly *Musca domestica*. The assay described here was designed to measure how flies execute specific, repeatable maneuvers requiring wingbeat amplitude modulation. Our data showed no significant differences in flight parameters between flies with intact gearboxes and those with ablated PWPs. Measurements of body trajectory, body angle, and wing angle remained comparable between the two groups. Additionally, the rates of yaw, pitch, and roll changes were similar, and wingbeat amplitudes during turns did not differ.

Within the constraints of the experimental design, we may therefore conclude that PWP is not necessary for wingbeat amplitude modulation and steering in flies. These conclusions challenge models that assign a central role to the PWP as a discrete “gearbox” mechanism for dipteran flight amplitude modulation. An alternative hypothesis proposes that instead of functioning as gears, the PWP grooves provide a textured surface that prevents the radial stop from slipping (Deora et al., 2015). In certain fly species, specific grooves may enhance friction, further reducing the likelihood of slippage between the RS and the PWP. This perspective implies that modes 1 and 2 are not distinct states; rather, amplitude modulation within these modes results from differential activity in the underlying steering muscles. Consequently, the gearbox mechanism may be less discrete and more dynamic than previously assumed.

## Acknowledgements

We acknowledge the Electron Microscopy Facility at NCBS, TIFR, Bengaluru for assistance with scanning electron microscopy. We thank Mr. Allan Joy for his help with insect collection, Dr Yeshwanth H.M. for help with insect identification, and Dr. Dinesh Natesan for his assistance with wing and body kinematics analysis. Funding for this research was provided by the Air Force Office of Scientific Research (AFOSR; grant numbers FA2386-11-1-4057 and FA9550-16-1-0155) and by NCBS, TIFR under project number 12-R&DTFR-5.04-0900.

## Supplementary information legends

**Movie S1. Top view of a *Musca domestica* performing a left turn maneuver.** The video was acquired at 4000 frames per second and is played back at 30 frames per second. The fly approaches from arm A of the L-box, rolls leftward, pitches upward, and then counter-rolls, effectively executing a banked turn.

